# *In Silico* Analysis of Non Synonymous Single Nucleotide Polymorphisms (nsSNPs) of SMPX Gene in Hearing Impairment

**DOI:** 10.1101/461764

**Authors:** Md. Arifuzzaman, Sarmistha Mitra, Amir Hamza, Raju Das, Nurul Absar, Raju Dash

## Abstract

**Background:** Mutations in SMPX gene can disrupt the normal activity of the SMPX protein which is involved in hearing process.

**Objective:** In this study, deleterious non-synonymous single nucleotide polymorphisms were isolated from the neutral variants by using several bioinformatics tools.

**Method:** Firstly, dbSNP database hosted by NCBI was used to retrieve the SNPs of SMPX gene, secondly, SIFT was used primarily to screen the damaging SNPs. Further, for validation PROVEAN, PredictSNP and PolyPhen 2 were used. I-Mutant 3 was utilized to analyze the protein stability change and MutPred predicted the molecular mechanism of protein stability change. Finally evolutionary conservation was done to study their conservancy by using ConSurf server.

**Results:** A total of 26 missense (0.6517%) and 3 nonsense variants (0.075%) were retrieved and among them 4 mutations were found deleterious by all the tools of this experiment and are also highly conserved according to ConSurf server. rs772775896, rs759552778, rs200892029 and rs1016314772 are the reference IDs of deleterious mutations where the substitutions are S71L, N19D, A29T and K54N. Loss of Ubiquitination, loss of methylation, loss of glycosylation, and loss of MoRF binding motifs are the root causes of protein stability change.

**Conclusion:** This is the first study regarding nsSNPs of SMPX gene where the most damaging SNPs were screened that are associated with the SMPX gene and can be used for further research to study their effect on protein structure and function, their dynamic behavior and how they actually affect protein’s flexibility.

## 1 INTRODUCTION

From the data of WHO (http://www.who.int), it is found that about 1 in 1000 newborns and a total of 360 million people worldwide have hearing loss (HL), which is a sensory disorder and extremely heterogeneous (1). HL can be classified as syndromic (SHL, 30%) and non-syndromic (NSHL, 70%) hearing loss and about 150 loci as well as 100 genes have been found to be associated with NSHL (Van Camp G, Smith RJH. http://hereditaryhearingloss.org) (2). X-linked non-syndromic hearing impairment is generally rare, with a percentage of around 1-5% and approximately 5% cases for pre-lingual male deafness (3). According to the recent study six NSHL loci have been identified and among them SMPX (DFNX4, OMIM 300066) is located in chromosome X at p.22.1 and consists of five exons where the first and the fifth exon being non-coding (4). SMPX gene encodes 88 amino acids and cloned firstly from muscle (4). The SMPX gene mainly conserved across mammalian species (5), and it has no known functional domains, and it is also involved primarily in inner ear development through the interaction with insulin-like growth factor 1 (IGF-1) (5, 6), integrins (α8β1) and Rac1 (7, 8). The SMPX protein exhibit the ability to protect the cells of the cochlea from the mechanical stress which is exerted during the process of hearing (9). Mutation in SMPX gene can cause hearing impairment with delayed detection and intervention and become more developing form of hearing loss (10, 11). Mutation can also cause significant damage to the inner ear cells and lead to progressive hearing loss (12). Several studies have already been conducted to identify the association between SMPX mutation and X-linked hearing loss in various families (13, 14). But no study was found to be carried out to analyze the involvement of single nucleotide polymorphisms (SNPs) of SMPX gene in X-linked hearing impairment. Single nucleotide polymorphisms are known as the most common types of variants occurring in human genome when a single nucleotide becomes substituted (15). Non-synonymous SNPs (nsSNPs) can cause 50% of the disease that are related to inheritance (16-19). From previous studies it was also revealed that, 60% of Mendelian disease caused by SNPs (20). SNPs can also disrupt the normal function of a protein and denature its structure by changing its stability, folding pattern and ligand binding site (21). For this reason, in this present experiment, we analyzed the involvement of nsSNPs of SMPX gene in X-linked hearing impairment by using various bioinformatics tools and computational databases.

## 2 METHODOLOGIES

### 2.1 Data retrieval

From the SNP database of National Center for Biotechnology Information (dbSNP) (http://www.ncbi.nlm.nih.gov/snp/), the corresponding datasets of SMPX gene were retrieved in accordance with their rsIDs which was considered as the first approach for data analysis (22).

### 2.2 Identification of damaging SNPs

For the isolation of deleterious SNPs and to analyze their impact on protein structure and function, seven major and widely accepted computational tools were used. As for theaim of this present study, only deleterious SNPs were considered rather than the neutral ones.

**SIFT** (Sorting intolerant from tolerant) (http://sift.jcvi.org) is a web based tool which was used to identify the damaging SNPs from the tolerated SNPs on the basis of the sequence homology (23-25). The prediction score based on a range of value, where a score ≤ 0.05 was considered as deleterious and a score ≥ 0.05 was considered as tolerated (26, 27). SIFT can be used by using SWISS-PROT, SWISS-PROT/TrEMBL, or NCBI’s non-redundant protein databases for SIFT to search (28, 29).

**PolyPhen2.0** (Polymorphism Phenotyping v2) (http://genetics.bwh.harvard.edu/pph2/uses) is another tool extensively used for prediction of functional impact of SNPs on protein structure and function (30). The input for this tool was the use of the respective protein sequence in FASTA format with the position of substitution and native as well as substituted amino acid, and it used Bayesian classifier method for prediction (31). Two prediction model was available such as HumDiv and HumVar (18). HumDiv mainly identified the less damaging SNPs whereas the HumVar classified the SNPs with extreme phenotypic effect on the basis of PSIC score (32). The probalistic score was ranging from 0 (neutral) to 1 (deleterious) and functional significance was categorized into benign (0.00–0.14), possibly damaging (0.15–0.84), and probably damaging (0.85–1) (33).

**PROVEAN** (http://provean.jcvi.org/index.php) was usedfor the prediction of changes in biological function of a protein due to amino acid substitution (34). This actually worked based on sequence clustering
and alignment based clustering score (35). For the generation of final PROVEAN score BLAST hits with more than 75% global sequence identity were clustered together and top 30 such clusters from a supporting sequence were averaged within and across clusters (36). The score of less than −2.5 was known as damaging and a score greater than −2.5 was considered as neutral (37).

**PredictSNP** (https://loschmidt.chemi.muni.cz/predictsnp/) is a very strong and the most recent tool for predicting damaging SNPs from the neutral SNPs, which accumulates various other bioinformatics tools and from the combined result the final PredictSNP result is generated which is very reliable and convenient (38). The most important tools which were used in this case are MAPP (39), nsSNPAnalyzer (40), PANTHER (41, 42), PhD-SNP (43), PolyPhen-1, PolyPhen-2 (30), SIFT (24) and SNAP (44). To be a damaging SNP, it must be validated by at least 3-4 tools among the all tools used in PredictSNP. The acquired results were provided together with annotations extracted from the Protein Mutant Database and the UniProt Database (38).

**I-mutant 3.0** (http://gpcr2.biocomp.unibo.it/cgi/predictors/I-Mutant3.0/I-Mutant3.0.cgi) is a support vector machine (SVM) based tool which was used to predict the protein stability change due to single nucleotide substitution (45). Input for this tool was the use of structure or sequence of the protein. We used the sequence based input for analysis. It predicted the protein stability change in three different classes such as, neutral mutation when the value of DDG was between −0.5 and 0.5 (−0.5 ≤ DDG ≤ 0.5 Kcal/mole), large decrease (< −0.5 kcal/mole) and large increase (> 0.5 kcal/mole) (43). The output file shows the predicted free energy change (DDG) value which was calculated from the folding Gibbs free energy of the mutated protein minus the unfolding free energy value of the native protein (43, 45).

**MutPred** (mutpred.mutdb.org/) was utilized for the analysis of protein’s structural and functional changes due to amino acid substitutions (46). It also predicted the molecular causes of deleterious mutations and also separated the damaging mutations from neutral mutations. It was trained using the deleterious mutations from the Human Gene Mutation Database (47) and the SIFT algorithm was used in this tool (24). The score range and prediction output can be, scores with *g* > 0.5 and *p* < 0.05 were referred to as actionable hypotheses, scores with *g* > 0.75 and *p* < 0.05 were referred to as confident hypotheses and scores with *g* > 0.75 and *p* < 0.01 were referred to as very confident hypotheses (46).

**ConSurf server** (consurf.tau.ac.il/) is a bioinformatics tool for the analysis of evolutionary conservation of an amino acid in a protein, DNA or RNA by using phylogenetic relationships among the homologous sequences. The phylogenetic tree was generated and evolutionary conservation analysis was done by using ConSurf server (48). Bayesian calculation method was used to calculate conservation score from protein sequence (49, 50). A conservation score between 1 and 4 was indicated as variable, whereas a score of 5–6 was intermediate, and a score between 7 to 9 indicated as conserved (51).

## 3 RESULTS

### 3.1 Dataset

A total of 10628 SNPs were found from preliminary search in dbSNP database of NCBI. Among them *Homo sapiens* comprises 3989 SNPs that means 37.5329% of total SNPs. Missense variants occupied 0.6517% with a number of 26 SNPs of total 3989 SNPs. 3 SNPs were nonsense which was only 0.075%. In intron, 3691 SNPs (92.53%), in 5’UTR region 21 SNPs (0.53%), in 3’UTR region 29 SNPs (0.727%), 16 (0.40%) SNPs as coding synonymous and 3 (0.075%) SNPs as frame shift variants were found. Only the non-synonymous missense and nonsesnse variants were then considered for further analysis.

### 3.2 Non-synonymous SNP analysis

After identifying 26 missense SNPs and 3 nonsense variants, we analyzed them by SIFT (**Table 1**) where 3 nonsense SNPs showed no substitution and one missense variant showed no result. So we did not consider the nonsense variants. As a result, we found the data of 25 missense SNPs. Out of 25; we got 15 SNPs as intolerated with a percentage of 60. SIFT analyzed the tolerance index (TI) as ≤ 0.05. SIFT requires the chromosome coordinates as input. The following rsIDs showed deleterious effect by scoring less than 0.05; rs150700337, rs200892029, rs369871483, rs376846054, rs745997977, rs747570937, rs756355143, rs759552778, rs761534067, rs763716855, rs765928656, rs770287561, rs772775896, rs779876319 and rs1016314772. Among the substitutions of those rsIDs; S71L, K62N, N19D, Q31L, E87G, R26W, P7T, A29T amino acid changes showed a score of 0. Out of 25 SNPs, 10 SNPs (40%) were found to be tolerated by scoring over 0.05 (**Table 6**).

**Table 1:**
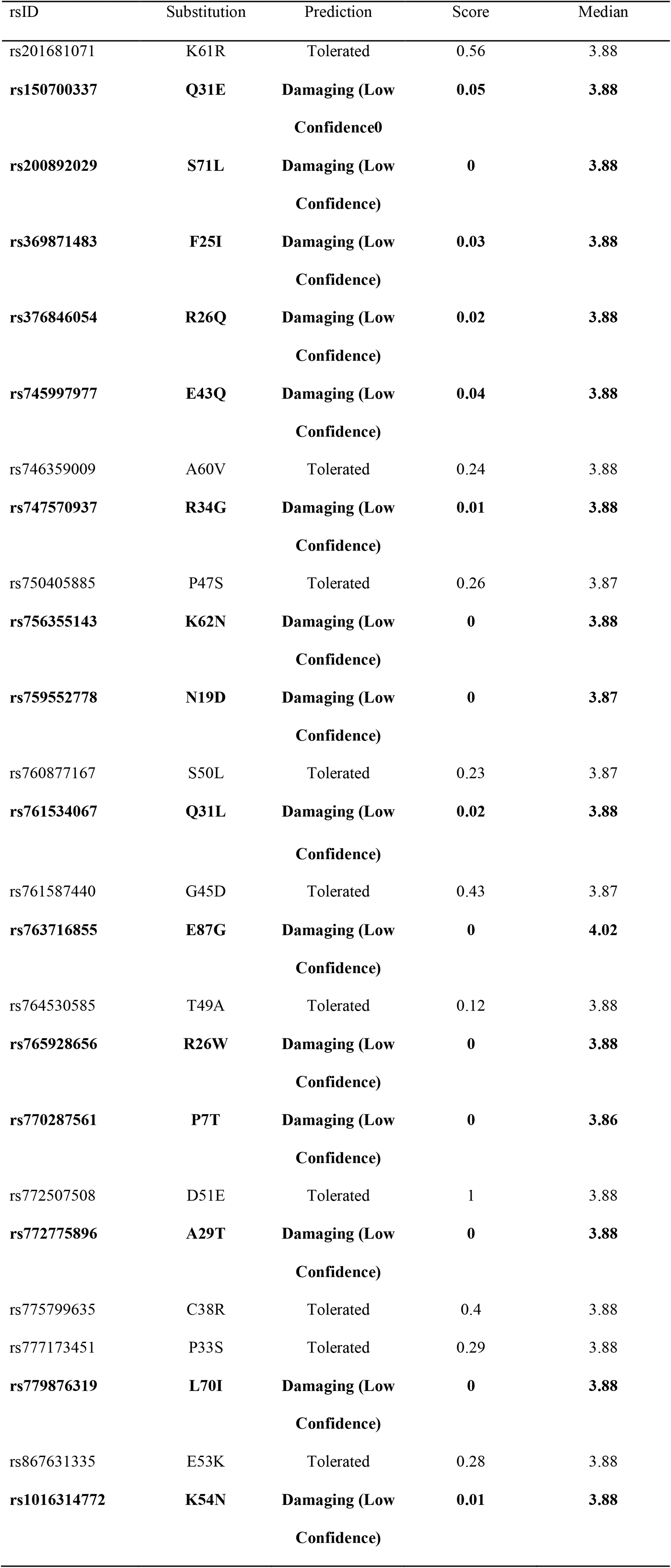
List of nsSNPs analyzed by SIFT (Damaging SNPs are indicated as bold)

**Table 2:** List of nsSNPs predicted by PolyPhen 2 (Damaging SNPs are indicated as bold, PRO= probably damaging, POS= possibly damaging, B= benign)

| rsID | Substitution | HumDIV | Value | HamVar | Value |
| --- | --- | --- | --- | --- | --- |
| rs201681071 | K61R | B | 0.004 | B | 0.003 |
| rs150700337 | Q31E | B | 0.341 | B | 0.06 |
| **rs200892029** | **S71L** | **PRO** | **1** | **PRO** | **0.992** |
| rs369871483 | F25I | POS | 0.705 | B | 0.271 |
| rs376846054 | R26Q | PRO | 0.998 | POS | 0.783 |
| rs745997977 | E43Q | B | 0.077 | B | 0.059 |
| rs746359009 | A60V | B | 0.198 | B | 0.061 |
| rs747570937 | R34G | B | 0.403 | B | 0.083 |
| rs750405885 | P47S | PRO | 0.959 | B | 0.362 |
| rs756355143 | K62N | B | 0.354 | B | 0.229 |
| **rs759552778** | **N19D** | **PRO** | **0.996** | **PRO** | **0.986** |
| rs760877167 | S50L | B | 0.049 | B | 0.016 |
| rs761534067 | Q31L | POS | 0.557 | B | 0.158 |
| rs761587440 | G45D | B | 0 | B | 0.001 |
| rs763716855 | E87G | B | 0.006 | B | 0.011 |
| rs764530585 | T49A | B | 0 | B | 0.001 |
| rs765928656 | R26W | PRO | 1 | POS | 0.88 |
| rs770287561 | P7T | B | 0.221 | B | 0.176 |
| rs772507508 | D51E | B | 0 | B | 0 |
| **rs772775896** | **A29T** | **PRO** | **1** | **PRO** | **0.996** |
| rs775799635 | C38R | B | 0.011 | B | 0.018 |
| rs777173451 | P33S | POS | 0.944 | POS | 0.451 |
| **rs779876319** | **L70I** | **PRO** | **0.996** | **PRO** | **0.992** |
| rs867631335 | E53K | B | 0.004 | B | 0.004 |
| **rs1016314772** | **K54N** | **PRO** | **0.999** | **PRO** | **0.994** |

**Table 3:** List of SNPs predicted by PROVEAN where cut off value = −2.5 (Deleterious SNPs are indicated as bold)

| Position | Wild residue | Mutant residue | Provean score | Provean prediction<br>(cutoff=-2.5) |
| --- | --- | --- | --- | --- |
| 61 | K | R | -1.89 | Neutral |
| 31 | Q | E | -1.72 | Neutral |
| **71** | **S** | **L** | **-5.83** | **Deleterious** |
| **25** | **F** | **I** | **-3** | **Deleterious** |
| **26** | **R** | **Q** | **-3.38** | **Deleterious** |
| 43 | E | Q | -2.22 | Neutral |
| **60** | **A** | **V** | **-3.27** | **Deleterious** |
| **34** | **R** | **G** | **-4.15** | **Deleterious** |
| **47** | **P** | **S** | **-3.38** | **Deleterious** |
| **62** | **K** | **N** | **-4.86** | **Deleterious** |
| **19** | **N** | **D** | **-4.15** | **Deleterious** |
| **50** | **S** | **L** | **-2.92** | **Deleterious** |
| **31** | **Q** | **L** | **-3.63** | **Deleterious** |
| 45 | G | D | -1.06 | Neutral |
| **87** | **E** | **G** | **-4.52** | **Deleterious** |
| 49 | T | A | -2.35 | Neutral |
| **26** | **R** | **W** | **-6.75** | **Deleterious** |
| **7** | **P** | **T** | **-6.56** | **Deleterious** |
| 51 | D | E | 0.08 | Neutral |
| **29** | **A** | **T** | **-3.5** | **Deleterious** |
| 38 | C | R | -2.35 | Neutral |
| **33** | **P** | **S** | **-4.16** | **Deleterious** |
| 70 | L | I | -2 | Neutral |
| **53** | **E** | **K** | **-2.73** | **Deleterious** |
| **54** | **K** | **N** | **-4.31** | **Deleterious** |

**Table 4:**
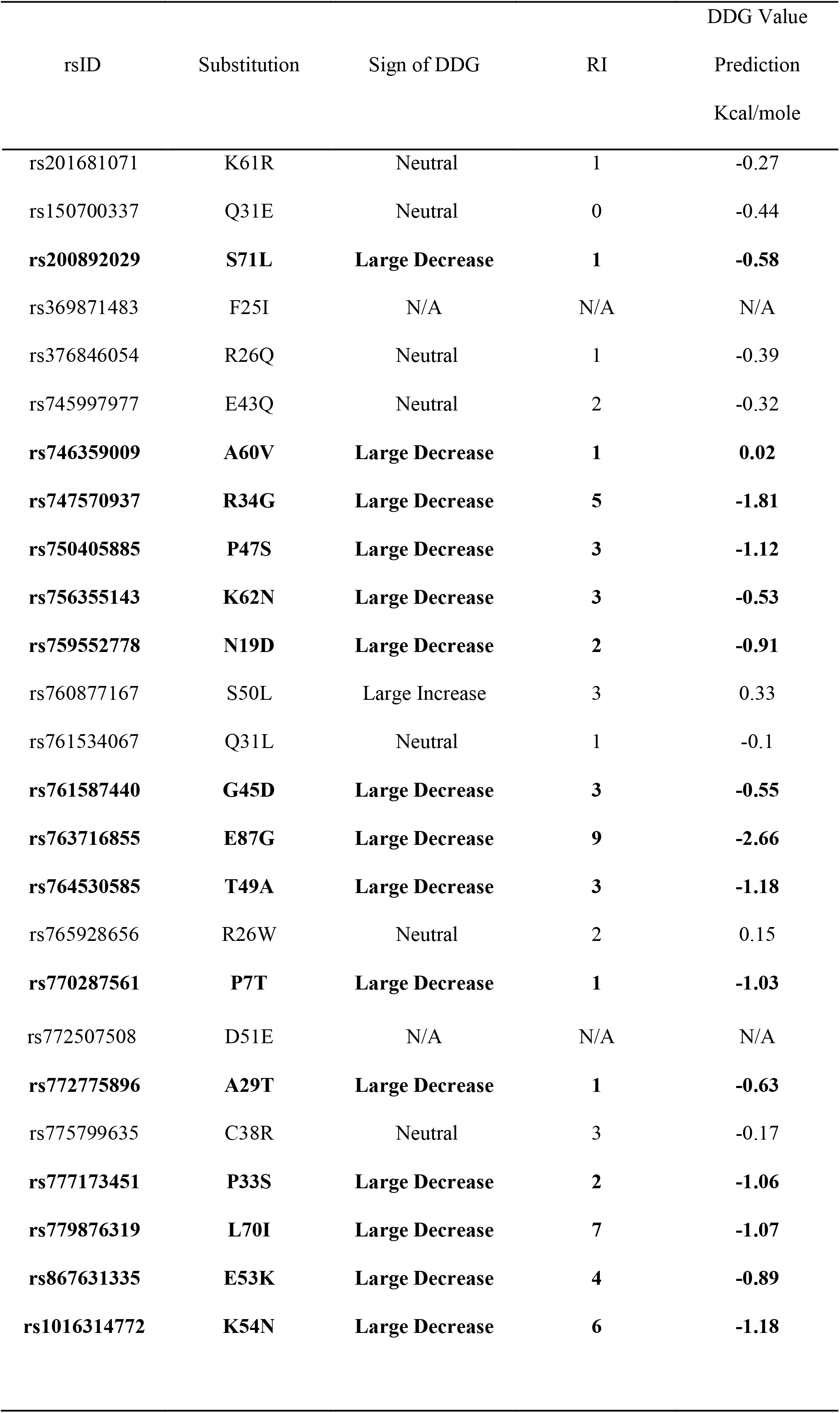
Protein stability change prediction by I-Mutant 3.0 (SNPs that are decreasing the stability of protein are indicated as bold)

| rsID | Substitution | Sign of DDG | RI | DDG Value |
| --- | --- | --- | --- | --- |
|  |  |  |  | Prediction |
|  |  |  |  | Kcal/mole |
| rs201681071 | K61R | Neutral | 1 | -0.27 |
| rs150700337 | Q31E | Neutral | 0 | -0.44 |
| **rs200892029** | **S71L** | **Large Decrease** | **1** | **-0.58** |
| rs369871483 | F25I | N/A | N/A | N/A |
| rs376846054 | R26Q | Neutral | 1 | -0.39 |
| rs745997977 | E43Q | Neutral | 2 | -0.32 |
| **rs746359009** | **A60V** | **Large Decrease** | **1** | **0.02** |
| **rs747570937** | **R34G** | **Large Decrease** | **5** | **-1.81** |
| **rs750405885** | **P47S** | **Large Decrease** | **3** | **-1.12** |
| **rs756355143** | **K62N** | **Large Decrease** | **3** | **-0.53** |
| **rs759552778** | **N19D** | **Large Decrease** | **2** | **-0.91** |
| rs760877167 | S50L | Large Increase | 3 | 0.33 |
| rs761534067 | Q31L | Neutral | 1 | -0.1 |
| **rs761587440** | **G45D** | **Large Decrease** | **3** | **-0.55** |
| **rs763716855** | **E87G** | **Large Decrease** | **9** | **-2.66** |
| **rs764530585** | **T49A** | **Large Decrease** | **3** | **-1.18** |
| rs765928656 | R26W | Neutral | 2 | 0.15 |
| **rs770287561** | **P7T** | **Large Decrease** | **1** | **-1.03** |
| rs772507508 | D51E | N/A | N/A | N/A |
| **rs772775896** | **A29T** | **Large Decrease** | **1** | **-0.63** |
| rs775799635 | C38R | Neutral | 3 | -0.17 |
| **rs777173451** | **P33S** | **Large Decrease** | **2** | **-1.06** |
| **rs779876319** | **L70I** | **Large Decrease** | **7** | **-1.07** |
| **rs867631335** | **E53K** | **Large Decrease** | **4** | **-0.89** |
| **rs1016314772** | **K54N** | **Large Decrease** | **6** | **-1.18** |

**Table 5:** List of SNPs predicted as deleterious by various tools used in this study.

| rsIDs | Substitutions | SIFT | PolyPhen 2 |  | PROVEAN | PredictSNP | I-Mutant 3 |
| --- | --- | --- | --- | --- | --- | --- | --- |
|  |  |  | HumDiv | HumVar |  |  |  |
| rs200892029 | S71L | Damaging<br>(Low<br>confidence) | PRO | PRO | Deleterious | Deleterious | Large<br>Decrease |
| rs759552778 | N19D | Damaging<br>(Low<br>confidence) | PRO | PRO | Deleterious | Deleterious | Large<br>Decrease |
| rs772775896 | A29T | Damaging<br>(Low<br>confidence) | PRO | PRO | Deleterious | Deleterious | Large<br>Decrease |
| rs1016314772 | K54N | Damaging<br>(Low<br>confidence) | PRO | PRO | Deleterious | Deleterious | Large<br>Decrease |

**Table 6:** The prediction results of SMPX gene obtained from SIFT, PolyPhen 2 and I-Mutant 3 of 25 nsSNPS

| Prediction | SIFT |  | PolyPhen |  | I-Mutant |  | PROVEAN |  | PredictSNP |  |
| --- | --- | --- | --- | --- | --- | --- | --- | --- | --- | --- |
|  |  |  | 2 |  | 3 |  |  |  |  |  |
|  | No. of | % | No. of | % | No. of | % | No. of SNPs | % | No. of SNPs | % |
|  | nsSNPs |  | nsSNPs |  | nsSNPs |  |  |  |  |  |
| Deleterious | 15 | 60 | 5 | 20 | 15 | 60 | 17 | 68 | 11 | 44 |
| Neutral | 10 | 40 | 20 | 80 | 10 | 40 | 8 | 32 | 14 | 56 |
| Total | 25 | 100 | 25 | 100 | 25 | 100 | 25 | 100 | 25 | 100 |

For the analysis of functional impact of those variants on protein structure and function PolyPhen 2 analysis was carried out (**Table 2**). From the results of PolyPhen 2 it can be observed that, 5 nsSNPs (20%) were deleterious and 20 nsSNPs (80%) were not deleterious (**Table 6**). The deleterious mutations showed both HumDiv and HumVar as probably damaging. The 5 variants were S71L, N19D, A29T, L70I and K54N. These 5 variants were also proved as damaging by SIFT.

PROVEAN is a clustering based method which was used for the validation of damaging SNPs which were selected and isolated from previous tools (**Table 3**). PROVEAN predicted 17 SNPs as deleterious (68%) by giving scores less than – 2.5 and 8 SNPs as neutral (32%) by scoring above – 2.5 (**Table 6**). PROVEAN also predicted SIFT score for accurate validation. The SNPs which were predicted as deleterious are S71L, F25I, R26Q, A60V, R34G, P47S, K62N, N19D, S50L, Q31L, E87G, R26W, P7T, A29T, P33S, E53K and K54N. Among these substitutions S71L, N19D, A29T and K54N were also predicted by PolyPhen 2.

PredictSNP tool was also used for further and final validation of nsSNPs that were also predicted and validated by SIFT, PolyPhen 2 and PROVEAN. According to this server, SMPX gene had 11 deleterious SNPs (44%) whereas 14 SNPs were defined as neutral (56%) (**Table 6**). The SNPs that were predicted as damaging are S71L, R26Q, K62N, N19D, G45D, T49A, R26W, P7T, A29T, L70I and K54N. Out of these S71L, N19D, A29T and K54N were also identified as deleterious by PROVEAN, PolyPhen 2 and SIFT. PredictSNP give prediction on the basis of 7 different tools which were mentioned in method section.

### 3.3 Prediction of Structural stability change due to mutations

I-Mutant 3.0 is a widely used tool for the prediction of stability change caused by the amino acid substitutions. I-Mutant 3.0 revealed that, 15 SNPs were decreasing the protein stability, 1 SNP was increasing the stability, 7 SNPs had no effect on stability of protein and 2 SNPs did not give any prediction, so this study considered them as neutral (**Table 6**). Here, 60% SNPs were deleterious and rest of them were 40%. The amino acid substitutions that were decreasing the stability are S71L, A60V, R34G, P47S, K62N, N19D, G45D, E87G, T49A, P7T, A29T, P33S, L70I, E53K and K54N. S50L was increasing the stability with the reliability index of 3 and DDG value with 0.33 (**Table 4**).

MutPred was used for the analysis of molecular mechanisms involved in decreasing the stability of protein by the SNPs. The SNPs which were showed deleterious effect by the previous tools such as SIFT, PolyPhen 2, PROVEAN, PredictSNP and I-Mutant 3 are accumulated in **Table 5** and we analyzed the molecular reason of protein stability change by those variants. Here, S71L showed that, it decreased protein stability by loss of loop (P=0.0512) and gain of helix (P = 0.062) which was a very confident hypothesis. Another substitution N19D caused loss of MoRF binding (P = 0.0441) and decreased protein stability was also very confident hypothesis. A29T also showed very confident hypothesis by gain of phosphorylation at A29 (P= 0.002). The last SNP, K54N, showed Loss of ubiquitination at K54 (P = 0.0011), loss of methylation at K54 (P= 0.0017) and loss of glycosylation at K54 (P= 0.0284) which was very confident hypothesis.

Finally evolutionary conservation analysis was performed by ConSurf server. It calculated the phylogenetic tree by using Bayesian classifier method of the homologous sequences. From the result of ConSurf server, it was found that, S71L, N19D, A29T and K54N were highly conserved **(Figure 1**).

**Figure 1:**
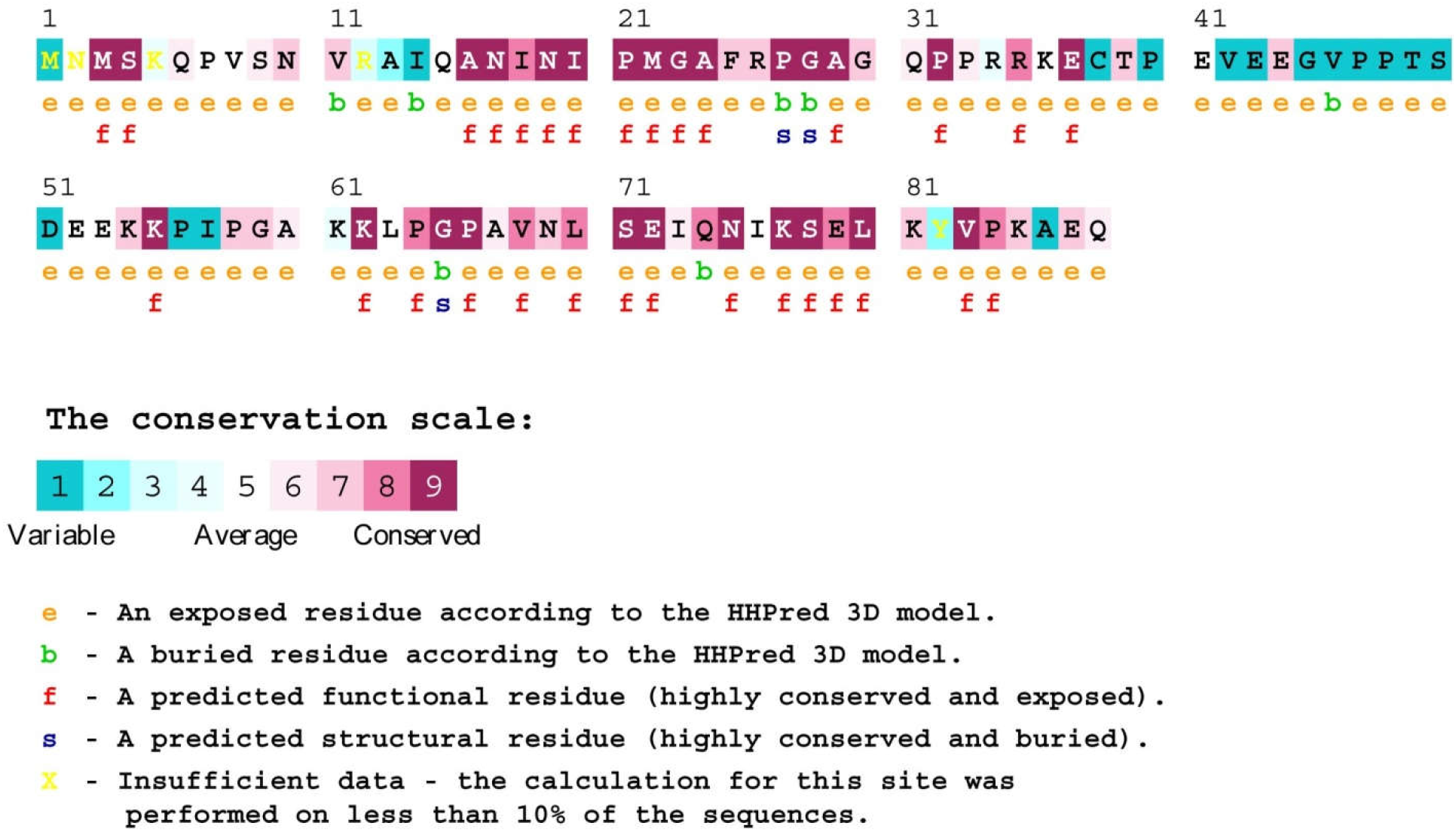
Evolutionary conservation analysis by ConSurf server

## 4 DISCUSSION

In recent days, identification of single nucleotide polymorphisms by *in silico* methods is very much popular to uncover the mutations that are involved in impairing protein structure and function. The SNPs of SMPX gene has not been considered till date to observe their functional and structural effects on protein. To address this issue, the present studt therefore used various *in silico* tools for the screening of deleterious SNPs from the neutral SNPs. In this study, various sequence based approaches were conducted because it is adventitious over structure based approaches. In structure based methods, it is limited to a known 3D structure; on the other hand, in sequence based approach it can be implemented to proteins with unknown 3D structure. In most studies, both sequence and structure based methods were used for validation of results and to make the results more reliable (52, 53). In our study, we first retrieved the SNPs from dbSNP database of NCBI for analysis. We found 26 SNPs as missense and 3 SNPs as nonsense. We used the missense variants for further analysis because we did not found any substitutions in case of nonsense variants. SIFT revealed that, 15 SNPs were damaging by scoring ≤ 0.05, 10 SNPs were tolerated by scoring ≥ 0.05 and 1 SNP did not give any result (**Table 1**). The SNPs that were found from SIFT are; Q31E, S71L, F25I, R26Q, E43Q, R34G, K62N, N19D, Q31L, E87G, R26W, P7T, A29T, L70I and K54N. For analysis of impact of SNPs on protein function, we used PolyPhen 2 server (**Table 2**). From the result of PolyPhen 2 it was found that, 5 SNPs were deleterious and 20 SNPs were neutral based on PSIC score. We considered both HumDiv and HumVar as probably damaging by scoring 0.85-1 of PSIC score. The deleterious SNPs were; S71L, N19D, A29T, L70I and K54N, which were also predicted as damaging by SIFT. To validate the results obtained from SIFT and PolyPhen 2, we considered PROVEAN (**Table 3**) and PredictSNP server. PROVEAN predicted 17 SNPs as deleterious by scoring < −2.5 and 8 SNPs as neutral by scoring > −2.5. S71L, F25I, R26Q, A60V, R34G, P47S, K62N, N19D, S50L, Q31L, E87G, R26W, P7T, A29T, P33S, E53K and K54N were predicted as damaging by PROVEAN and among them S71L, N19D, A29T and K54N were also predicted as damaging by SIFT and PolyPhen 2. PredictSNP combines several tools under the same platform such as; MAPP, nsSNPanalyzer, PolyPhen 1, PolyPhen 2, SIFT, PANTHER and SNAP. According to the result of PredictSNP, 11 SNPs were indicated as deleterious whereas 14 SNPs as neutral. The following SNPs are damaging; S71L, R26Q, K62N, N19D, G45D, T49A, R26W, P7T, A29T, L70I and K54N. Out of these S71L, N19D, A29T and K54N were also identified as deleterious by PROVEAN, PolyPhen 2 and SIFT. Afterward, we used I-Mutant 3 for the analysis of protein stability change due to mutations. 15 SNPs were known to decrease the stability of protein such as S71L, A60V, R34G, P47S, K62N, N19D, G45D, E87G, T49A, P7T, A29T, P33S, L70I, E53K and K54N. 1 SNP was increasing the stability of protein and 7 SNPs had no effect on protein function (**Table 4**). Finally, the molecular mechanisms behind the loss of function of the protein due to variants were analyzed by using MutPred server. The SNPs that were predicted as deleterious by all the tools used in this study (**Table 5**) for MutPred analysis. The MutPred server revealed that, S71L decreased protein stability by loss of loop (P=0.0512) and gain of helix (P = 0.062) which was a very confident hypothesis. Another substitution N19D caused loss of MoRF binding (P = 0.0441) and decrease protein stability which was also very confident hypothesis. A29T also showed very confident hypothesis by gain of phosphorylation at A29 (P= 0.002). The last SNP K54N showed loss of ubiquitination at K54 (P = 0.0011), loss of methylation at K54 (P= 0.0017) and loss of glycosylation at K54 (P= 0.0284) which was also very confident hypothesis. The SNPs that were conserved across the evolutionary perspective are more important than who were not conserved because they are structurally and functionally important for the protein (49). For this reason, ConSurf server was used for evolutionary conservation analysis **(Figure 1**). This analysis showed that; S71L, N19D, A29T and K54N were highly conserved.

## 5 CONCLUSION

Our study overviewd critically all of the deleterious and neutral SNPs that were involved in SMPX gene. Overall, from the analysis, four SNPs have been found to be mostly deleterious predicted by all tools used in this experiment and may affect hearing and cause hearing impairment. S71L, N19D, A29T and K54N were the single nucleotide polymorphisms identified by SIFT, PolyPhen 2, PROVEAN, PredictSNP, I-Mutant 3 and MutPred. These SNPs were also highly conserved according to the conservancy analysis by ConSurf server. Further experiment is required to study their dynamic behavior, how they affect protein structure and function by increasing or decreasing the flexibility of the protein.

